# Root xylem in three woody angiosperm species is not more vulnerable to embolism than stem xylem

**DOI:** 10.1101/2020.03.30.017343

**Authors:** Min Wu, Ya Zhang, Thais Oya, Carmen Regina Marcati, Luciano Pereira, Steven Jansen

## Abstract

**Aims:** Since plants are compartmentalised organisms, failure of their hydraulic transport system could differ between organs. We test here whether xylem tissue of stems and roots differ in their drought-induced embolism resistance, and whether intact roots are equally resistant to embolism than root segments.

**Methods:** Embolism resistance of stem and root xylem was measured based on the pneumatic technique for *Acer campestre*, *A. pseudoplatanus* and *Corylus avellana*, comparing also intact roots and root segments of *C. avellana*. Moreover, we compared anatomical features such as interconduit pit membrane between roots and stems.

**Results:** We found a higher embolism resistance for roots than stems, although a significant difference was only found for *A. pseudoplatanus.* Interconduit pit membrane thickness was similar for both organs of the two *Acer* species, but pit membranes were thicker in roots than stems of *C. avellana*. Also, embolism resistance of an intact root network was similar to thick root segments for *C. avellana*.

**Conclusion:** Our observations show that root xylem is not more vulnerable to embolism than stem xylem, although more species need to be studied to test if this finding can be generalised. We also demonstrated that the pneumatic method can be applied to non-terminal plant samples.

## Introduction

The plant hydraulic system is known to form a soil-plant-atmosphere continuum (Taiz and Zeiger 1998). Plants transport water in the xylem vascular system, pulling water from the soil through roots, stems, and leaves (Dixon 1896). The continuous water column in the xylem vascular system can be interrupted by embolism (i.e. the entry of large gas bubbles in conduits) (Jansen and Schenk 2015; Lens et al., 2011). Although embolism can occur in the xylem of any plant organ, an interesting question is whether or not different organs are equally vulnerable to embolism. The answer to this question is especially relevant because of the compartmentalised nature of plants, which affects not only anatomy, but also functional processes such as transport and defence systems (Morris et al. 2016a, 2019; Schenk et al. 2008).

The “hydraulic vulnerability segmentation hypothesis” (HVSH) suggests that distal organs (e.g. leaves and roots) are less resistant to xylem embolism than the proximal organs (e.g. stems and trunks) by means of providing hydraulic safety and protecting vital, meristematic tissues on proximal organs from dehydration (Johnson et al. 2016; Tyree and Ewers 1991; Zhu et al. 2016). This hypothesis is supported by some studies on angiosperms (e.g. *Acer pseudoplatanus*, *Fagus sylvatica*, *Juglans regia* × *nigra* and *Juglans regia*; Cochard et al. 2002; Hochberg et al. 2016; Losso et al. 2019; Tyree et al. 1993) and gymnosperms (e.g. *Cupressus sempervirens* and *Pinus halepensis*; Domec et al. 2006; Froux et al. 2005; Sperry and Ikeda 1997). Based on these studies, leaves and roots are found to be less resistant to embolism than branches and trunks (Creek et al. 2018; Johnson et al. 2016). However, other studies show clear evidence against the HVSH. For instance, similar xylem embolism resistance between leaves and branches was found in *Allocasuarina verticillata*, *Betula pendula*, *Eucalyptus pulchella*, *Liriodendron tulipifera*, *Melaleuca pustulata* and *Pinus pinaster* (Bouche et al. 2016; Klepsch et al. 2018; Smith-Martin et al. 2020). Similar xylem embolism resistance was also found in xylem from roots, stem and leaves of tomato plants (Skelton et al. 2017). Moreover, embolism resistance of roots was found to be higher than stems instead of lower for *Juniperus ashei*, *Quercus fusiformi*s, *Q. sinuata*, *Olea europaea*, and several poplar species (Hukin et al. 2005; McElrone et al. 2004; Rodriguez-Dominguez et al. 2018). The discrepancy between studies may be due to variation among species, potential differences in habitats and growth patterns, and/or methodological differences in the protocol applied (Choat et al. 2016; Lamarque et al. 2018; Wheeler et al. 2013).

Variation in xylem embolism resistance is typically reflected in xylem anatomy (Choat et al. 2008; Jansen et al. 2009; Tyree and Zimmermann 2002). The characteristics of interconduit pits, especially the pit membrane thickness, are suggested to be one of the major determinants of xylem embolism resistance (Lazzarin et al. 2016; Li et al. 2016a; Prendin et al. 2018; Tissier et al. 2004). There is strong and convincing evidence that drought-induced embolism is determined by pit membranes, which control embolism spreading between neighbouring conduits (Kaack et al. 2019). Interestingly, interconduit pit membranes showed a similar thickness in leaves and branches of *Liriodendron tulipifera* and *Laurus nobilis*, but were found to be thicker in leaves than in branches of *Betula pendula* (Klepsch et al. 2018). Quantitative variation in interconduit pit membrane thickness among various organs within a single tree of *Acer pseudoplatanus* showed considerable variation (Kotowska et al. 2020), but it is unknown whether pit membrane thickness may determine differences in xylem embolism resistance among organs within a single plant. Moreover, xylem anatomical features are known to show various quantitative differences between stems and roots, which reflect also their different mechanical properties (Plavcová et al. 2019).

This study investigates xylem embolism resistance in stems and roots of three angiosperm species (*Acer campestre*, *A. pseudoplatanus*, and *Corylus avellana*) based on the manual pneumatic method (Pereira et al. 2016; Zhang et al. 2018). Moreover, anatomical observations of stem and root xylem were carried out to test for potential differences in xylem anatomy between these organs. Our main objectives are 1) to test whether or not stems and roots of the species selected differ in their xylem embolism resistance, and whether or not this difference is associated with xylem anatomical characteristics; 2) to examine whether embolism resistance is similar between intact root systems and non-terminal root segments. Finally, an automated pneumatron device was applied to intact roots of *C. avellana* and compared with the manual pneumatic approach. These tests are essential to develop straightforward protocols to estimate embolism vulnerability in roots, allowing comparisons among plant organs.

## Materials and methods

### Plant material

All measurements were performed on three common temperate angiosperm trees: *Acer campestre*, *A. pseudoplatanus*, and *Corylus avellana*. These species were selected based on availability of plant material and access to their roots. Five to ten saplings, which were ca. 0.75 to 1 m tall and 4 to 5 years old, were grown in 3 L pots at the Botanical Garden of Ulm University (48°25’ 9.84’’N, 9°57’ 59.7 6’’E). The measurements were conducted between April (after leaf flushing) and October 2018. The saplings were growing in a mixture of organic soil / sand / loam peat / turf (5:2:2:1, v/v/v/v) and watered on a regular basis during the growing season. The stem and the intact root system of each sapling were collected for our measurements.

Moreover, five saplings of *C. avellana*, which were ca. 2 m in height and 4 to 5 years old, were selected for their root segments. These sapling were growing in the forest of the Botanical Garden of Ulm University.

The pneumatic method (Pereira et al. 2016; Zhang et al. 2018) was applied to potted saplings of three species, and forest saplings of *C. avellana* (Fig. 1). This method measures the amount of gas extracted from xylem tissue over time, while the sample is left to dry at room temperature (Bittencourt et al. 2018; Fig. 2). Based on a striking correlation between the amount of gas extracted in pneumatic measurements and measurements on the loss of hydraulic conductivity for more than 20 tropical and temperate angiosperm species (Pereira et al. 2016; Zhang et al. 2018), there is very good empirical evidence that the pneumatic method quantifies embolism resistance (Barros et al. 2019; Brum et al. 2019; Lima et al. 2018; Oliveira et al. 2019).

**Fig. 1.**
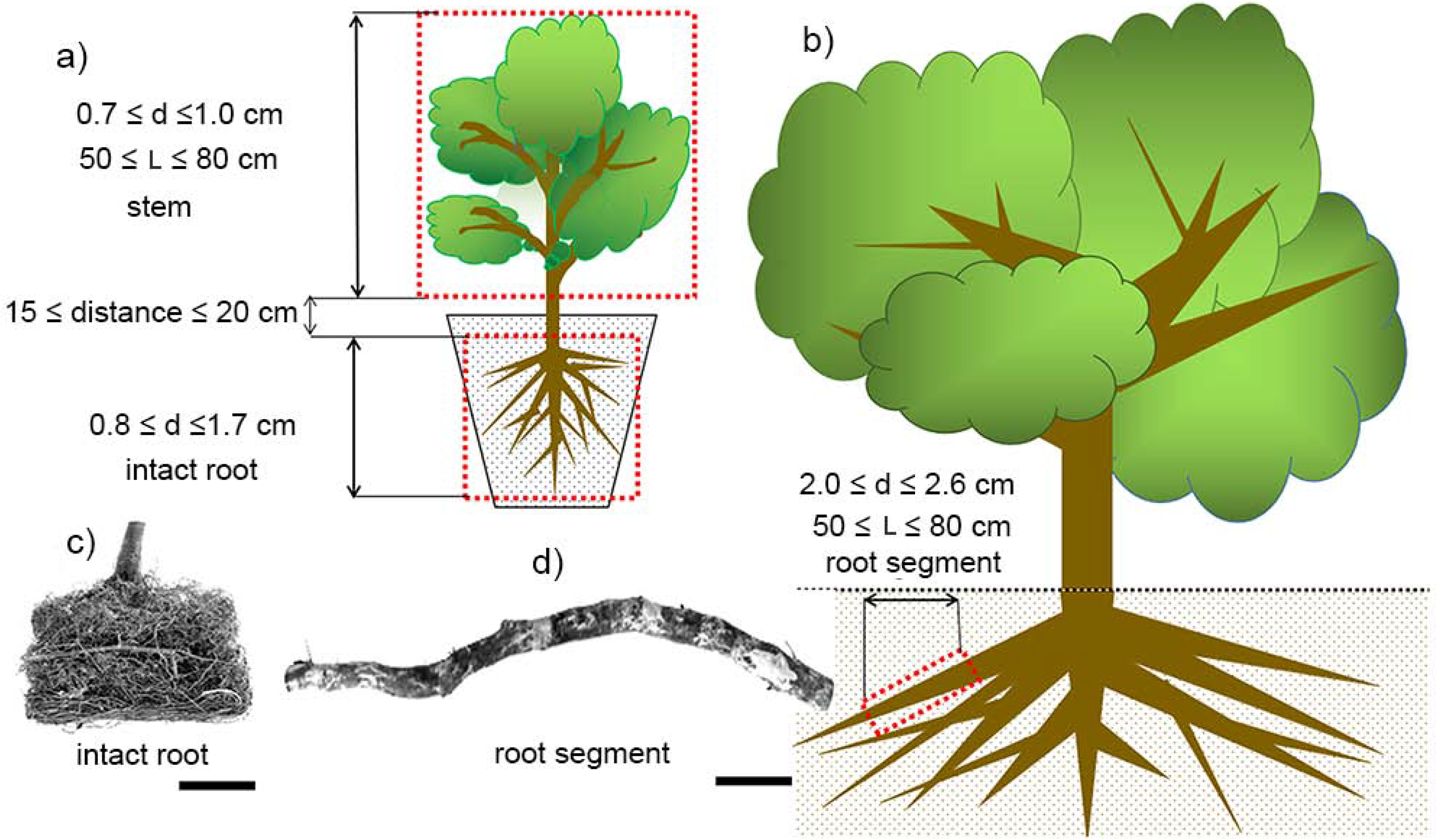
Scheme for sample preparation. For each potted seedling, the stem and the intact root were separated at the stem-root junction, where a 15-20 cm transition part was removed. The stem was then further trimmed to a length of 50-80 cm, with a stem diameter of 0.7-1.0 cm at the proximal end, and the diameter of the intact root was 0.8-1.7 cm (a). For saplings of *C. avellana*, a root segment of 50-80 cm long with a diameter of 2.0-2.6 cm at the proximal end was dug out manually (b). d = sample diameter, L = sample length. Pictures show an intact root network of *A. pseudoplatanus* (c), and a mature root segment of *C. avellana* (d). Scale bars in (c) and (d) = 15 mm

**Fig. 2.**
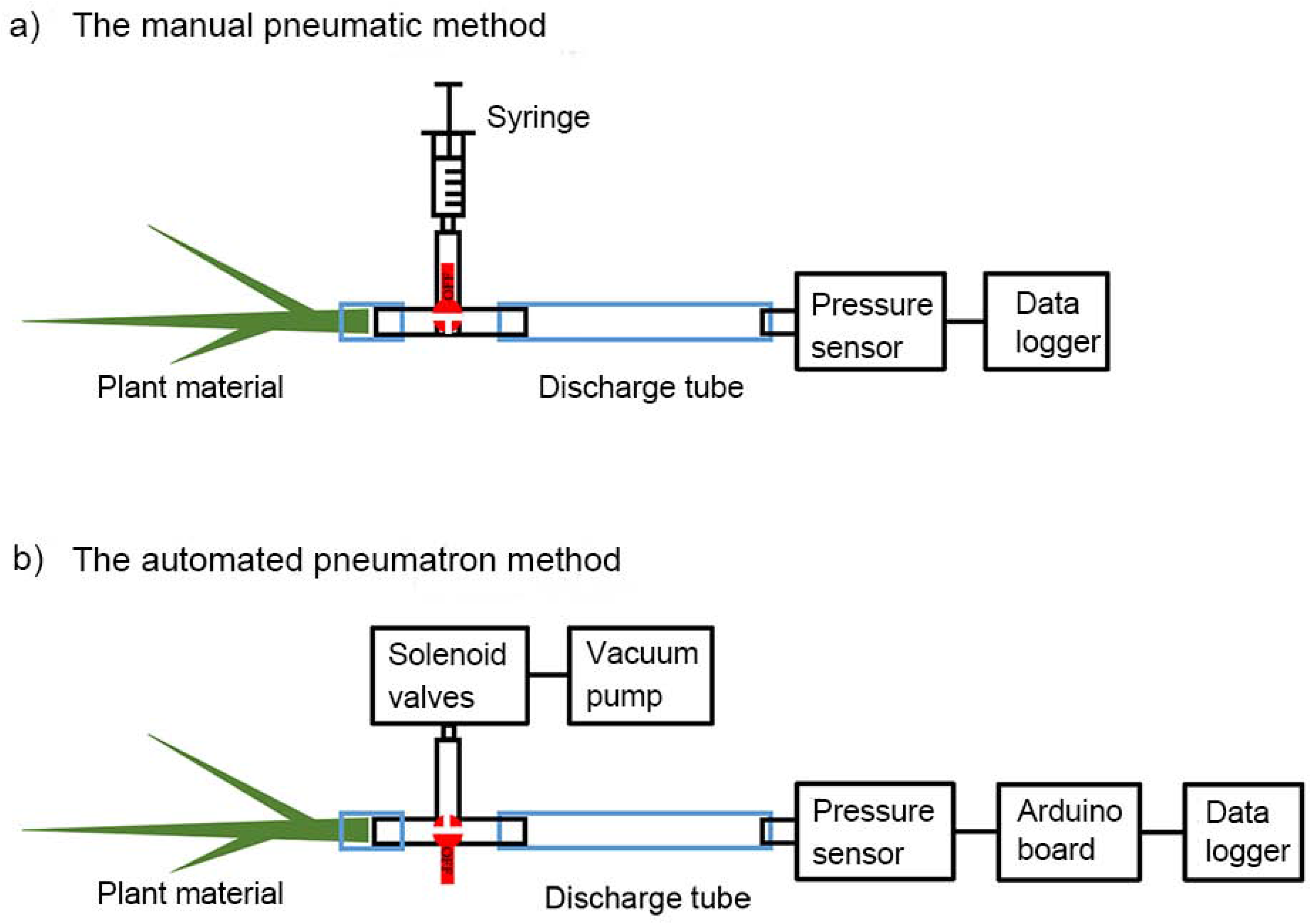
Schematic diagrams of the manual pneumatic (a) and the automated pneumatron method (b). Both approaches measure the amount of air discharged from plant samples over time. (a) A plant sample (e.g. stem, intact root, or root segment) was connected to a three-way stopcock, which was linked to a syringe and a rigid discharge tube. The syringe acted as a vacuum generator and the tube as a vacuum reservoir. Increasing pressure in the discharge tube were recorded by a pressure sensor (PX26-015GV) and saved in a data logger (CR850). Before opening the sample-vacuum reservoir pathway, a pressure of 40 kPa was applied in the discharge tube by carefully pulling the syringe. Then, the sample-vacuum reservoir pathway was opened and air from the sample was discharged into the rigid tube with a known volume. This diagram was modified from Pereira et al. (2016). (b) The automated pneumatron method was generally similar to the setup of the manual pneumatic approach. The syringe was replaced by a vacuum pump together with solenoid valves. Increasing pressure in the discharge tube was recorded by the same pressure sensor and automatically saved in a data logger shield. An Arduino board was added to take pneumatic measurements at a constant time interval (e.g. each 15 min)

Xylem vulnerability curves were conducted for (1) stems (terminal branch ends), (2) intact root networks of potted saplings, and (3) root segments of *C. avellana* saplings growing in the forest. For each species, five potted saplings were wrapped with moist paper towels in the early morning, and transported in sealed black plastic bags to the laboratory. Saplings were taken out of the pots and carefully washed with tap water to clean the roots. The stem and the entire root network of each sapling were separated under water from the stem-root junction. Both stems and root networks were kept under water to rehydrate for 1 h. Then, the stem was recut under water at the proximal end and shortened into 50-80 cm long branches, which had a diameter of 0.7-1.0 cm at the cut end. The root network was kept intact, including many fine, highly curled roots, and had a root diameter of 0.8-1.7 cm at the cut surface. The stem-root junction that was cut off was ca. 15-20 cm long.

For the five saplings of *C. avellana* that were collected in the forest, one root segment per sapling was carefully dug out in the early morning in August 2018. Segments of 50-80 cm long and 2.0-2.6 cm thick were wrapped in wet paper tissue, enclosed in a black plastic bag, and transferred to the laboratory. Root segments were kept under water to rehydrate for 1 h, carefully trimmed with a razor blade, and the proximal end was connected to the pneumatic apparatus for embolism resistance measurements. The other (distal) end of the root segments was sealed with super glue (Loctite 431).

### Vulnerability to embolism

#### The manual pneumatic approach

The pneumatic apparatus (Fig. 2a) consists of a syringe (0.06 L) as a vacuum generator, a rigid tube (0.0082 L) as sample-vacuum reservoir, and a pressure sensor (PX26-015GV, Omega Engineering, NJ, USA) and a data logger (CR850, Campbell Scientific, Logan, USA). A pressure of 40 kPa was applied in the tube via the syringe and kept for 2 min so that any potential leakage (0-1 kPa) could be recorded. Then, the proximal end of hydrated stems or roots were connected to the tube by opening a three-way stopcock to the sample-vacuum reservoir pathway and the initial pressure (*P*_i_, kPa) in the tube was recorded immediately. After 2 min, the final pressure (*P*_f_, kPa) was also recorded. This procedure was repeated several times until the samples were severely dehydrated, which occurred at a xylem water potential (Ψ, MPa) of −9.5 MPa for stems, and −9.0 MPa for roots. An important observation of the pneumatic method is that the final pressure value *P*_f_ at the beginning of a measurement differs only slightly from the initial pressure *P*_i_ when there is hardly any embolism, but rises to a maximum difference when most vessels are embolised. This observation makes clear that in between pneumatic measurements the amount of gas inside a xylem sample is restored to its original concentration by equilibration under atmospheric pressure. In between pneumatic measurements, samples were left drying on a bench and bagged up in a black plastic bag for 60 min to obtain a water potential equilibrium within the sample. The drying time was 20 min for the first measurements, and one to four hours for the last measurements.

The xylem water potential of each sample was determined with a pressure chamber (PMS 1000, PMS Instruments, Albany, USA) immediately after conducting a pneumatic measurement. We took the mean value of the water potential of two leaves cut from a stem, or two lateral roots from an intact root network. The cut tissue was immediately sealed with super glue (Loctite 431) to avoid air leakage from the cut surface. For the root segments of *C. avellana*, which showed little to no side roots, the xylem water potential was determined with a stem psychrometer (PSY1, ICT International, Armidale, NSW, Australia), which was attached near the middle of the root segment.

Based on the ideal gas law, the amount of moles of air discharged (Δ*n*, mol) for each pneumatic measurement was computed as follows:

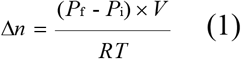

where *V* was the volume of the sample-vacuum reservoir (0.0082 L in our pneumatic apparatus), *R* was the gas constant (8.314 kPa L mol^−1^ K^−1^), and *T* was the room temperature in the laboratory. Then, the volume of the air discharged (AD, μl) was calculated based on the ideal gas law by transforming Δ*n* to an equivalent volume of air at atmospheric pressure (Patm, 94 kPa at 618 m, the altitude of Ulm University). Any potential leakage from the apparatus over 2 min was then subtracted from AD (the volume of air discharged).

The percentage of air discharged (PAD, %) was calculated as follows:

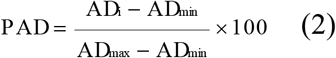

where AD_i_ was the volume of air discharged for each measurement, AD_min_ was the minimum volume of air discharged when the sample was fresh, and AD_max_ was the maximum volume of air discharged when the sample was at its lowest xylem water potential.

Vulnerability curves were then generated by plotting PAD against their corresponding water potential (Ψ) values with the following equation (Pammenter and Vander Willigen 1998):

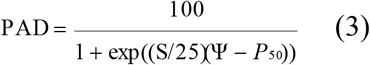

where S showed the slope of the curve and *P*_50_ was the xylem water potential at 50% of air discharged from samples. *P*_88_, which represented the xylem water potential at 88% of air discharged from samples, was calculated using the following equation (Domec and Gartner 2001):

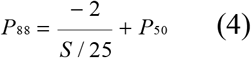

#### The pneumatron method

A pneumatron device (Fig. 2b; Pereira et al. 2020) was applied to five potted saplings of *C. avellana* to obtain xylem vulnerability curves of intact roots. The pneumatron is an automated pneumatic apparatus, including a microcontroller (Arduino Uno board; Adafruit Industries, NY, USA), a pressure sensor (PX26-015GV, Omega Engineering, NJ, USA), a mini-vacuum pump (DQB380-FB2, Dyx, Shenzhen, China), two mini-solenoid valves (DSF2-A, Dyx, Shenzhen, China), and a data logger shield (Adafruit Industries, NY, USA).

Briefly, a vacuum (absolute pressure = 45 kPa) created by the pump was applied to a rigid tube that was connected with the proximal end of an intact root. The pressure in the tube was recorded every 0.5 s and saved on a SD card by a data logger module. After 2.5 min, the measurements were finished. The next measurement started automatically after 12.5 min. The intact root was put on a lab bench and gradually desiccating during the measurements. The pneumatron measurements were stopped manually when the intact root showed severe dehydration. The xylem water potential of an intact root was determined by measuring the water potential of two lateral roots using the pressure chamber. The xylem water potential was monitored every 20 min at the beginning of dehydration, and every 1-4 hour as dehydration proceeded. The volume of air discharged (AD) and the percentage of air discharged (PAD) were calculated in a similar way as described above. Finally, vulnerability curves were built to obtain *P*_50_ and *P*_88_ values.

### Anatomical measurements

Light microscopy (LM), scanning (SEM) and transmission electron microscopy (TEM) were conducted at Ulm University. For LM and SEM, one stem and one lateral woody root segment from a single potted sapling per species were collected after conducting embolism resistance measurements. Five individuals were included for each species. TEM observations were based on one fresh (i.e. not drought stressed) stem xylem and one fresh lateral root from a single sapling per species. The TEM sample was not taken from the branches used for pneumatic measurements, because pit membranes would then be aspirated and partly or completely shrunken.

#### Light microscopy (LM)

To determine the conduit diameter (D, μm), small blocks (ca. 10 × 10 × 10 mm) of stem and lateral root segments were softened with a 25% (v/v) glycerin for 8-10 h. Then, transverse sections of 10 μm thick were obtained from each block with a sliding microtome (Schenkung Dapples Mikrol). Sections were stained with a 1% safranin solution for 5 min, and then washed twice with distilled water for 30 s. Then, they were dehydrated through an ethanol series (50%, 70% and 96% EtOH) for 2 min each. The stained sections were transferred to a microscope slide and embedded with Neo-Mount (Merck Millipore). The slides were dried in an oven at 60℃ for 15h and observed under a light microscope. Photographs of transverse sections were taken with a digital camera (Zeiss Axio Zoom V16, Göttingen, German) and the conduit diameter was manually measured using ImageJ (version 1.48v, National Institutes of Health, Bethesda, MD, USA) (Schindelin et al. 2012) based on 250 counts for each organ per species (Scholz et al. 2013a).

#### Scanning electron microscopy (SEM)

Xylem segments (5-10 mm in length) of stems and roots were split with a sharp blade to expose radial surfaces. These samples were air-dried at room temperature for one week, fixed to aluminium stubs and sputter-coated with gold-palladium (FL-9496 Balzers, Fürstentum Liechtenstein) for 2 min. Finally, these samples were examined under a SEM (Phenom-XL-0067-L, Netherlands) at an accelerating voltage of 5 kV.

SEM images were used for measuring the bordered pit membrane area (A_PM_, μm^2^), pit aperture area (A_PA_, μm^2^) and interconduit pitfield fraction (F_PF_) (the ratio of interconduit surface area occupied by interconduit pits to the total interconduit wall area) with a minimum of 50 pits included for each organ per species. The pit aperture fraction (F_PA_) was calculated as the ratio of pit aperture area (A_PA_) to pit membrane area (A_PM_) (Lens et al. 2011). The pit density (P_D_, No. per 100 μm^2^) was defined as the number of pits per 100 μm^2^ interconduit area, and was measured for a minimum of 10 conduits for each organ per species (Scholz et al. 2013a; Zhang et al. 2017). Special care was taken to distinguish vessel-vessel and vessel-tracheid pits from vessel-parenchyma pits. The latter could be identified based on their spatial arrangement.

#### Transmission electron microscopy (TEM)

Fresh blocks (1 to 2 mm^3^) of xylem from the current growth ring were cut under distilled water, and immediately stored in a standard fixative solution (2.5% glutaraldehyde, 0.1 mol phosphate, 1% sucrose, pH 7.3) in a fridge overnight. Samples were washed in 0.1 M phosphate buffer, post-fixated with 2% buffered osmium tetroxide for 2 h, and stained with uranyl acetate for 30 min at 37℃. Then, samples were dehydrated with a gradual ethanol series (30%, 50%, 70%, 90% and 100%), immersed in 1.2-propylenoxide (CAS260 Nr. 75-56-9, Fontenay-sous-Bois cedex), and gradually embedded in Epon resin (Sigma-Aldrich, Steinheim, Germany), which was polymerized at 60℃ over 48 h. Semi-thin sections (ca. 500 nm thick) were prepared from embedded samples with an ultramicrotome (Leica Ultracut UCT, Leica Microsystems, Vienna, Austria), stained with 0.5% toluidine blue in 0.1 M phosphate buffer, and mounted on slides with Eukitt. Ultra-thin sections (70-90 nm thick) were cut using a diamond knife and mounted on 300 mesh hexagonal copper grids (Agar Scientific, Stansted, U.K.). Observations were conducted with a JEOL1400 TEM (JEOL, Tokyo, Japan) at 120 kV accelerating voltage, and images were taken with a MegaView III camera (Soft Imaging System, Münster, Germany).

Since pit membranes showed a rather homogeneous thickness in TEM images, pit membrane thickness (T_PM_, nm) was determined as the mean value of three measurements, i.e. at opposite sides near the pit membrane annulus and in the centre (Zhang et al. 2017). At least 10 interconduit pit membranes were measured using ImageJ (Schindelin et al. 2012) for each organ per species. Interconduit wall thickness (T_CW_, μm) was defined as the double wall thickness of two neighbouring conduits and measured for at least 50 replicates based on semi-thin sections for each organ per species. In most cases, these conduits included vessels, but since the imperforate nature of some conduits could not be determined in transverse sections with absolute certainty, we used the term interconduit wall thickness to include both vessels and tracheids.

### Statistics

SPSS software (version 21, IBMCorp. Armonk, NewYork) was used for statistical analyses. Comparison of xylem embolism resistance and anatomical features between stems and roots at the intraspecific level were conducted using independent sample t-tests after testing for normal distribution (Sharpiro-Wilk test) and homogeneity of variance (Levene-test). Similar t-tests were applied to determine differences in xylem embolism resistance traits between the manual pneumatic method and the automated pneumatron method for intact roots of *C. avellana*, and between intact root networks and root segments of *C. avellana*. All figures were made with SigmaPlot 12.5 (Systat Software Inc., Erkrath, Germany).

## Results

### Variation in stem and root vulnerability to xylem embolism

Vulnerability curves of stems and intact root networks of the three species were obtained with the manual pneumatic method (Fig. 3). All vulnerability curves showed a sigmoidal shape. The 95% confidence bands of the stem and root curves showed considerable overlap for *A. campestre* and *C. avellana* (Fig. 3a and c), but not for *A. pseudoplatanus* (Fig. 3b).

**Fig. 3.**
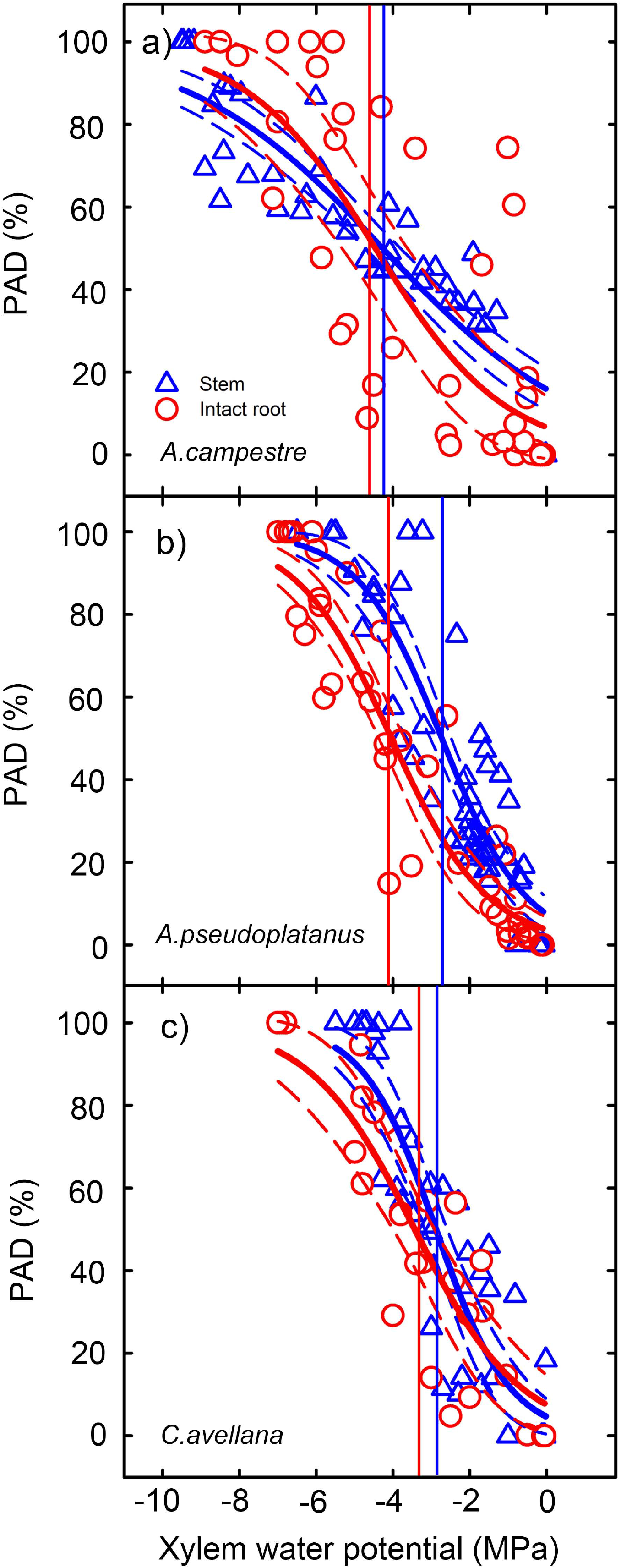
Xylem vulnerability curves of stems and intact roots from three angiosperm tree species (a, b, c) based on the manual pneumatic method. Blue triangles and blue lines represent data and curves for stems, and red circles and red lines represent data and curves for intact root networks. Solid lines show the regression curves and dash lines show 95% confidence bands. Solid vertical lines represent xylem water potential at 50% embolism (*P*_50_). Vulnerability curves were based on five individuals for each species. PAD = the percentage of air discharged

Xylem in roots showed more negative *P*_50_ values than stems (Fig. 4a). In *A. campestre*, the mean *P*_50_ value in root xylem was −4.86 ± 0.84 MPa, but was not significantly different (t(6) = 0.702, *P* = 0.509) from the stem xylem *P*_50_ (−4.21 ± 0.38 MPa) (Fig. 4a; Tables S1 and S2). Similarly, no significant difference in *P*_50_ (t(8) = 1.882, *P* = 0.097) was found between roots and stems of *C. avellana*, which showed a mean value of −3.52 ± 0.33 MPa and −2.77 ± 0.22 MPa, respectively (Fig. 4a; Tables S1and S3). For *A. pseudoplatanus*, however, the mean *P*_50_ value of root xylem was −4.17 ± 0.31 MPa, which was significantly more negative (t(4) = 3.924, *P* = 0.004) than the stem xylem *P*_50_ (−2.63 ± 0.24 MPa; Fig. 4a; Tables S1 and S2).

**Fig. 4.**
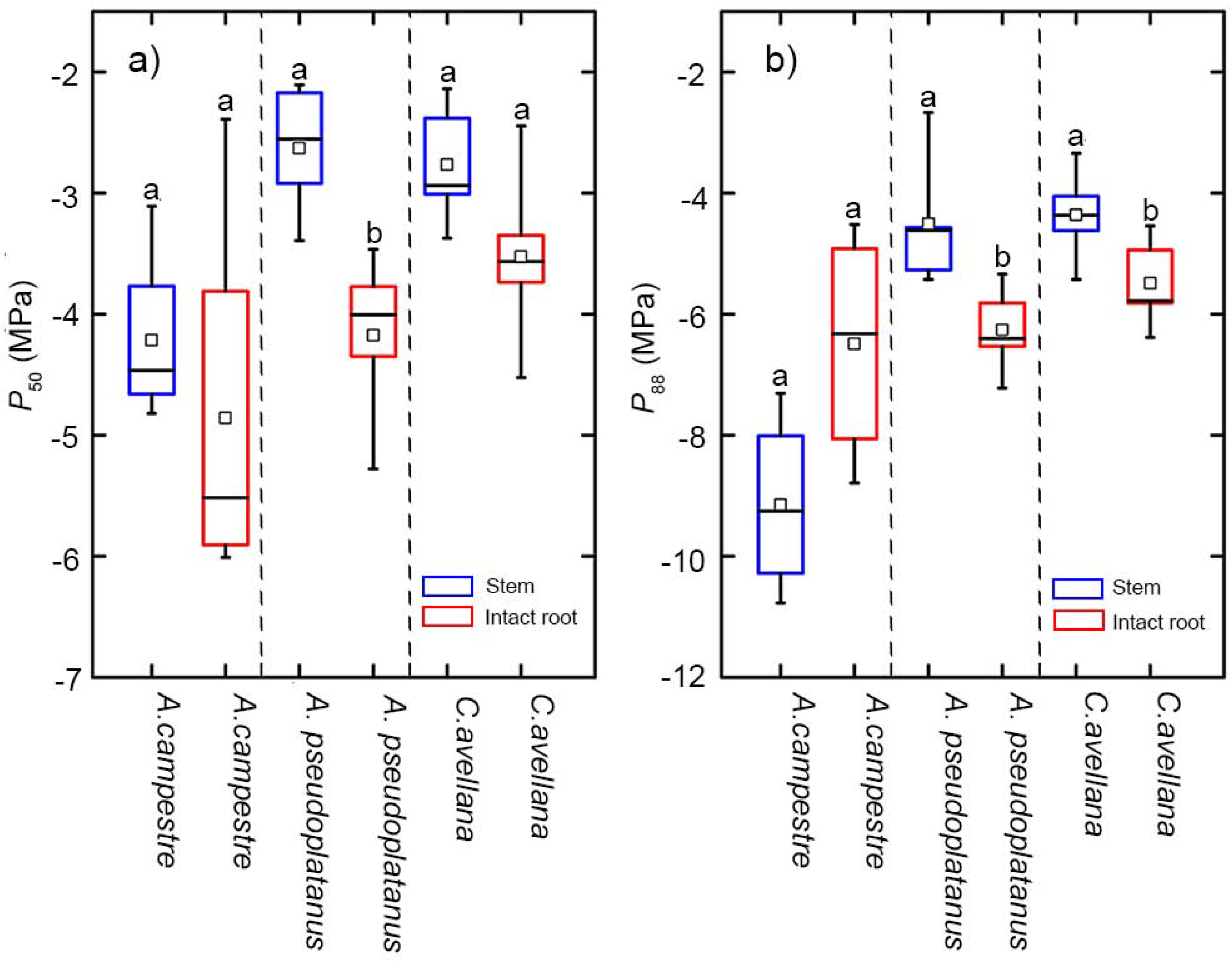
Comparisons of the *P*_50_ (a) and *P*_88_ (b) values (n = 5) between stems and intact root networks of three angiosperm tree species based on the manual pneumatic method. Different small letters indicate significant differences. Data of stems and roots were present in blue and red, respectively. Box plots show the median (horizontal line inside the box), average (square inside the box), 90th percentile (upper bar), 10th percentile (lower bar), 75th percentile (upper box line), and 25th percentile (lower box line)

A comparison of *P*_88_ values between xylem of stems and roots showed no significant difference (t(6) = −2.182, *P* = 0.072) for *A. campestre*, with the stem *P*_88_ (−9.15 ± 0.74 MPa) being more negative than the root *P*_88_ (−6.49 ± 0.97 MPa) (Fig. 4b; Tables S1 and S2). The opposite was shown for *A. pseudoplatanus* and *C. avellana*, with root *P*_88_ values (−6.26 ± 0.32 MPa and −5.49 ± 0.33 MPa, respectively) being significantly more negative than stem *P*_88_ values (−4.51 ± 0.49 MPa and −4.36 ± 0.34 MPa, respectively) (Fig. 4b; Tables S1 and S2).

### Variation in root vulnerability to xylem embolism in *C. avellana*

There was no significant difference (t(6) = −0.225, *P* = 0.830) in embolism resistance between intact root networks from potted saplings and thick root segments from forest saplings, with a mean *P*_50_ value of −3.52 ± 0.33 MPa and −3.41 ± 0.30 MPa, respectively (Fig. 5a, b and d; Tables S1 and S3). Similarly, no significant difference in *P*_88_ values (t(6) = 0.123, *P* = 0.906) was found between intact roots and thick roots segments of *C. avellana* (Fig. 5e; Tables S1 and S3).

**Fig. 5.**
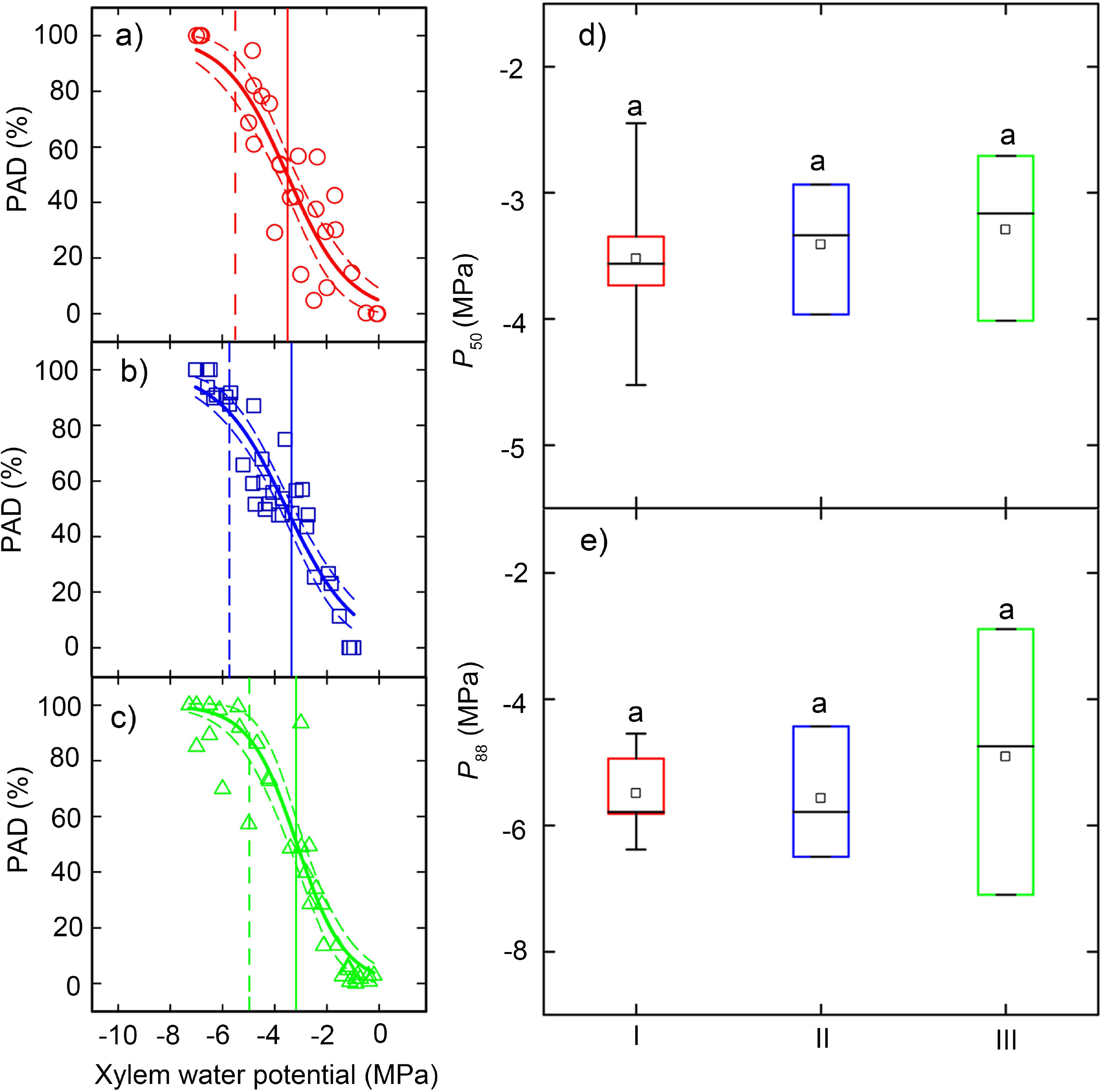
Root xylem vulnerability curves of *C. avellana*, with intact root networks based on the manual approach (a), thick root segments based on the manual pneumatic method (b), and intact root networks based on the automated pneumatic method (c). Solid lines show the fitting curves and the short dash lines represent the 95% confidence bands. Solid and dash vertical lines represent *P*_50_ and *P*_88_ values, which are shown in the boxplots d and e, respectively. No difference in *P*_50_ (d) and *P*_88_ (e) values (n = 3 or 5) were found. I = application of the manual method to intact root networks (red), II = the manual pneumatic method applied to thick roots segments (blue), and III = application of the automated pneumatron to intact root networks (green). PAD = the percentage of air discharged. Box plots show the median (horizontal line inside the box), average (square inside the box), 90th percentile (upper bar), 10th percentile (lower bar), 75th percentile (upper box line), and 25th percentile (lower box line)

No significant difference (t(6) = −0.435, *P* = 0.679) was found between *P*_50_ values based on the manual approach and the automated pneumatron method (Fig. 5a, c and d; Tables S1 and S3), with *P*_50_ values of −3.52 ± 0.33 MPa and −3.30 ± 0.38 MPa, respectively. A similar trend was shown in *P*_88_ values, with no significant difference (t(6) = 0.142, *P* = 0.579) based on these two methods (Fig. 5e; Tables S1 and S3).

### Conduit anatomy

Conduit diameter (D) was significantly wider in roots than in stems of the species studied (*P* < 0.001) (Table 1). Significant differences were found in interconduit cell wall thickness (T_CW_) between stem xylem and root xylem of *A. campestre*, *A. pseudoplatanus*, and *C. avellana* (*P <* 0.01), with the mean interconduit cell wall thickness being higher in stems than in roots (Table 1).

**Table 1.** Wood anatomical features related to conduit and pit characteristics for stem and root xylem in three angiosperm tree species.

| Species | Organ | D<br>($\mu\text{m}$) | A<sub>PM</sub><br>($\mu\text{m}^2$) | A<sub>PA</sub><br>($\mu\text{m}^2$) | P<sub>D</sub><br>(No. per 100 $\mu\text{m}^2$) | F<sub>PA</sub><br>(%) | F<sub>PF</sub><br>(%) | T<sub>PM</sub><br>(nm) | T<sub>CW</sub><br>($\mu\text{m}$) |
| --- | --- | --- | --- | --- | --- | --- | --- | --- | --- |
| *A. campestre* | Stem | 19.6 ± 0.4 | 26.6 ± 0.9 | 2.0 ± 0.2 | 2.2 ± 0.2 | 8.0 ± 0.8 | 63.1 ± 1.4 | 235 ± 12 | 3.1 ± 0.1 |
| | Root | 26.2 ± 0.6* | 24.9 ± 1.1 | 1.4 ± 0.1* | 2.1 ± 0.2 | 6.0 ± 0.4 | 53.9 ± 5.8 | 248 ± 11 | 2.7 ± 0.1* |
| *A. pseudoplatanus* | Stem | 24.4 ± 0.5 | 29.2 ± 1.1 | 2.0 ± 0.1 | 1.7 ± 0.2 | 7.3 ± 0.5 | 55.4 ± 6.7 | 256 ± 14 | 3.2 ± 0.1 |
| | Root | 27.7 ± 0.6* | 28.2 ± 1.5 | 2.3 ± 0.2 | 1.7 ± 0.2 | 10.0 ± 1.0* | 54.9 ± 4.7 | 266 ± 12 | 2.9 ± 0.1* |
| *C. avellana* | Stem | 25.8 ± 0.4 | 23.9 ± 1.8 | 3.1 ± 0.2 | 3.2 ± 0.3 | 16.2 ± 1.5 | 59.4 ± 2.0 | 241 ± 10 | 4.0 ± 0.1 |
| | Root | 29.7 ± 0.6* | 26.3 ± 1.6 | 3.0 ± 0.1 | 3.1 ± 0.6 | 14.3 ± 1.0 | 57.6 ± 4.4 | 304 ± 14* | 3.2 ± 0.1* |

D = conduit diameter, A<sub>PM</sub> = bordered pit area, A<sub>PA</sub> = pit aperture area, P<sub>D</sub> = pit density, F<sub>PA</sub> = pit aperture fraction, F<sub>PF</sub> = interconduit pit field fraction, T<sub>PM</sub> = interconduit pit membrane thickness, and T<sub>CW</sub> = interconduit double wall thickness of conduit. Values are means ± SE, n = 5. \* = statistical significance ( $P < 0.05$ ) between stems and roots.

### Bordered pit characteristics

There were both similarities and differences in the bordered pit structure between stem and root xylem (Table 1; Fig. 6). The pit border surface area (A_PM_) showed no significant difference between stem and root xylem (*P* > 0.05, Table 1; Fig. 6). Pit borders had a slightly larger surface area in stems than roots for *A. campestre* and *A. pseudoplatanus*, but the opposite was found for *C. avellana* (Table 1; Fig. 6). The pit aperture surface area (A_PA_) showed no difference between stems and roots (*P* > 0.05), except for *A. campestre* (t(122) = −2.945, *P* = 0.005). For *A. campestre*, pit aperture areas were larger in the stem than in the root (Table 1).

**Fig. 6.**
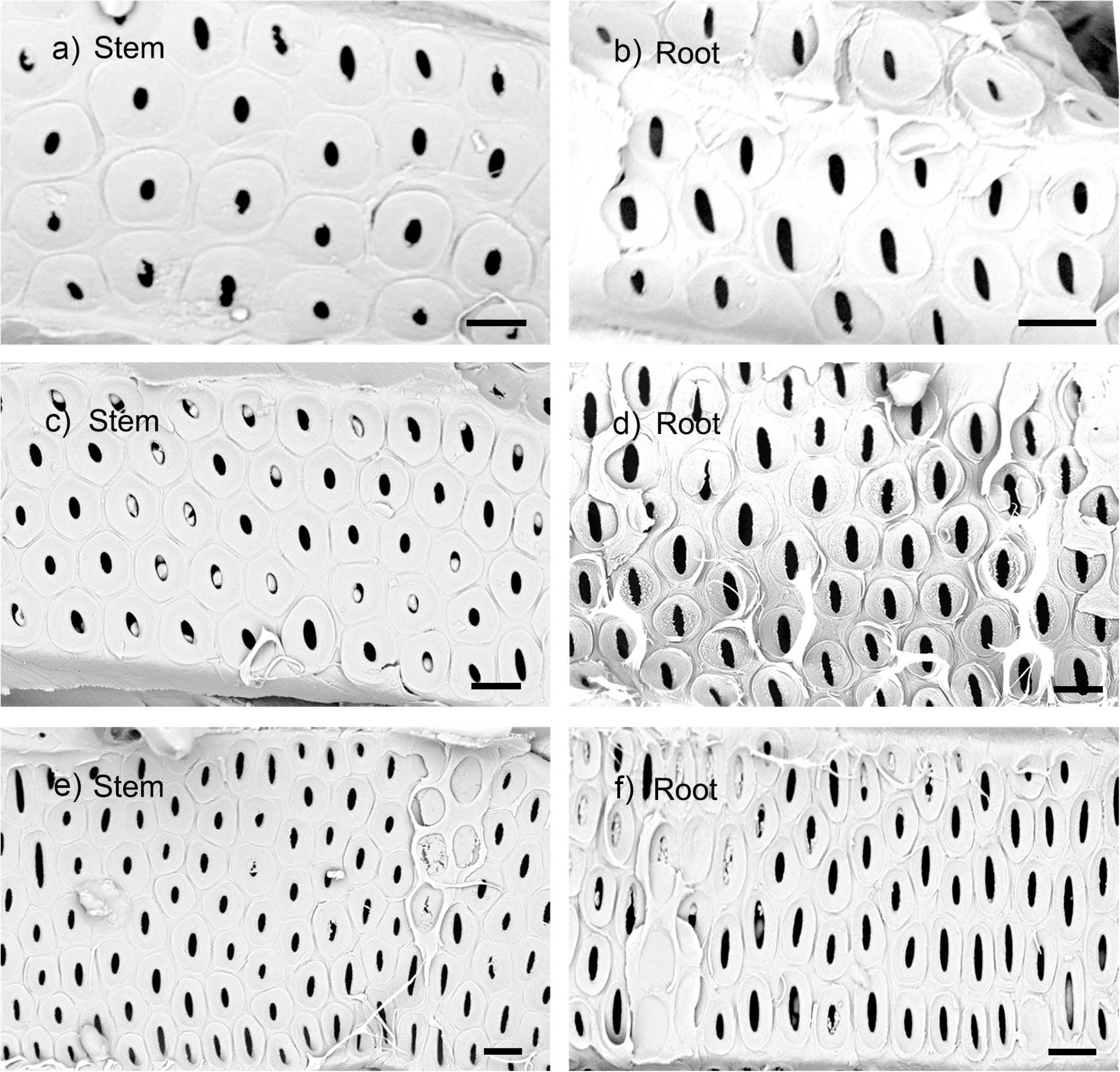
SEM images of interconduit pits in stem (a, c, e) and root (b, c, f) xylem of *A. campestre* (a, b), *A. pseudoplatanus* (c, d), and *C. avellana* (e, f). The axial orientation of the conduits is in a horizontal position for all images. All scale bars = 5 μm

The ratio between the pit aperture surface area and pit membrane surface area (F_PA_) was slightly higher in stems than roots for *A. campestre* and *C. avellana* (*P* > 0.05, Table 1). The pit aperture area represented about 7% and 15% of the total pit border area for stems and roots of *A. campestre* and *C. avellana*, respectively. This ratio was significantly lower in stems than roots for *A. pseudoplatanus* (t(107) = 2.281, *P* = 0.025), with pit apertures occupying 7% of the pit border area in the stem, and 10% in root xylem (Table 1).

There was no significant difference in the mean pit density (P_D_) and the interconduit pit field fraction (F_PF_) between stem and root xylem (*P* > 0.05, Table 1). Mean values of pit density and interconduit pit field fraction were higher in stem than in root xylem, except for pit density values in *A. pseudoplatanus* (1.73 ± 0.17 and 1.74 ± 0.15 per 100 μm^2^ of interconduit area for stem and root xylem, respectively).

TEM observations of intercoduit pit membranes of fresh samples showed considerably darker (i.e. more electron dense) particles in roots than in stems for *A. pseudoplatanus* and *C. avellana*, but not for *A. campestre* (Fig. 7). Most interconduit pit membranes showed a homogeneous appearance in electron density (Fig. 7b, c and e). Pit membranes with both transparent and electron dense parts were observed in roots of *A. pseudoplatanus* and *C. avellana*, with darker particles generally at the outermost layers of the pit membrane (Fig. 7d and f). The interconduit pit membranes were slightly thicker in root xylem than stem xylem for the three species studied (Table 1). The difference in pit membrane thickness, however, was only significant between stem and root xylem for *C. avellana* (t(67) = −3.704, *P* < 0.001) (Table 1).

**Fig. 7.**
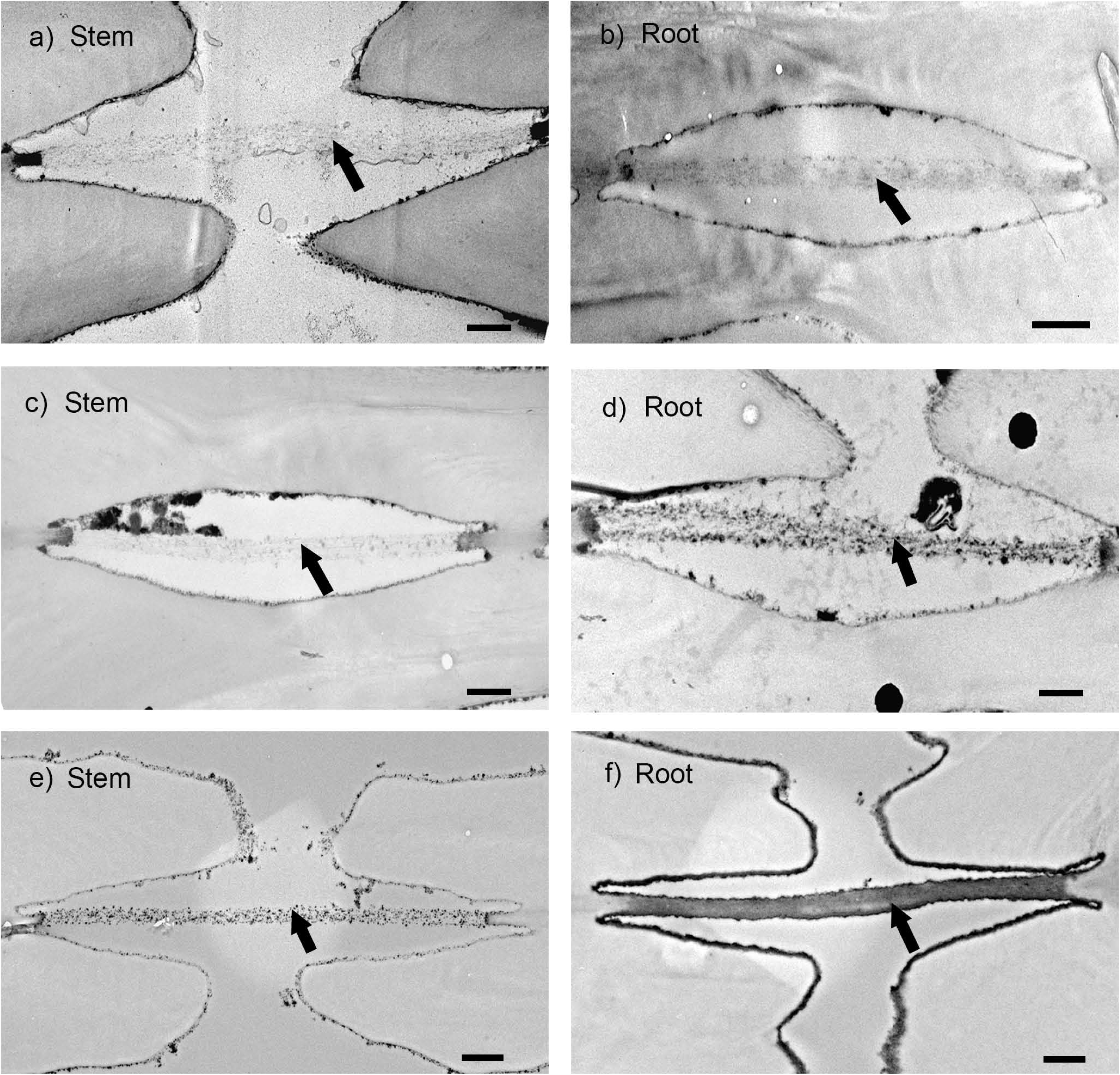
TEM images of interconduit pit membranes in stem (a, c, e) and root (b, d, f) xylem of *A. campestre* (a, b), *A. pseudoplatanus* (c, d), and *C. avellana* (e, f). All scale bars = 0.5 μm. Black arrows show the pit membranes

## Discussion

Embolism resistance of intact roots was found to be similar to stems for *A. campestre* and *C. avellana*, but higher than stems for *A. pseudoplatanus*. This finding is in line with previous studies that show no universal occurrence of the hydraulic vulnerability segmentation hypothesis (Hukin et al. 2005; McElrone et al. 2004; Rodriguez-Dominguez et al. 2018; Skelton et al. 2017), and raises some important points. Firstly, whether or not embolism occurs more quickly in a distant organ than in a proximal organ depends on the potential difference in xylem water potential that these organs experience in the field. Although we did not take xylem water potential measurements in the field, stems have a typically more negative water potential than roots (Jackson et al. 2000). Therefore, even without any pronounced difference in embolism resistance, hydraulic safety margins (defined as the difference between the xylem water potential experience and the *P*_50_ value) might be lower for stems than for roots.

Secondly, the discrepancy between our results and earlier studies that report more vulnerable xylem in roots than in stems (Domec et al. 2006; Pratt et al. 2007, 2015) could at least be partly affected by the different methods applied to measure xylem embolism resistance. Low root embolism resistance, for instance, has especially been found for angiosperms and conifers based on the air-injection method with double-ended cavitation chambers (Domec et al. 2006; Froux et al. 2005; Johnson et al. 2016; Martínez-Vilalta et al. 2002; Sperry and Ikeda 1997). This method may underestimate embolism resistance due to effervescence (i.e. the escape of gas bubbles from xylem sap when stem segments are exposed to reduced gas pressure according to Henry’s law, resulting in artificial embolism) or other methodological concerns (Choat et al. 2010; Torres-Ruiz et al. 2014, 2015; Yin and Cai 2018). High vulnerability to embolism may also be found for centrifuge methods, especially when vessels are at least half as long as the rotor diameter (Du et al. 2018; Hacke et al. 2015; Lamarque et al. 2018; Sperry et al. 2012; Torres-Ruiz et al. 2017; Wang et al. 2014). Observations based on X-ray tomography and the optical method suggest that embolism resistance of intact roots is similar to stems (Losso et al. 2019; Skelton et al. 2017), or even higher than stems (Rodriguez-Dominguez et al. 2018). The *P*_50_ values based on the pneumatic method for stems of *A. campestre* and *A. pseudoplatanus* are 1 to 2.2 MPa more negative than *P*_50_ values obtained with the air-injection method (Tissier et al. 2004). However, our *P*_50_ values for stems of the three species differed only by 0.4 to 0.7 MPa to those reported by Li et al. (2016b), which were based on the cavitron method using samples from the same population of trees. Moreover, our mean *P_50_* value for stems of *A. pseudoplatanus* (−2.63 MPa) is also similar to the −2.51 MPa *P*_50_ based on micro-CT observations (Losso et al. 2019). These similarities are interesting and confirm earlier comparison of the pneumatic method with the bench dehydration approach (Pereira et al. 2016; Zhang et al. 2018).

One major advantage of the pneumatic method is that its measurements are based on the kinetics of gas diffusion from embolised, non-cut open conduits via interconduit pit membranes to the discharge tube. However, extraction of gas from alternative sources (e.g. extraxylary tissue or intercellular air spaces in xylem) is much slower (Sorz and Hietz 2006), and can therefore be ignored. This also means that the pneumatic method does not measure any change in root hydraulic conductivity, which may be strongly affected by extraxylary tissue prior to embolism formation in xylem conduits (Cuneo et al. 2016; Rodriguez-Dominguez and Brodribb 2020).

When both xylem and extraxylary conductivity were considered (Creek et al. 2018), roots were found to be more vulnerable than stems. However, our study showed an opposite trend based on the pneumatic measurements where the extraxylary tissue were not measured. *P*_50_ methods that are unable to distinguish extraxylary conductivity from xylem conductivity could overestimate xylem embolism resistance.

The finding that root xylem was not more vulnerable to embolism than stem xylem based on our pneumatic measurements was further supported by data on pit membrane thickness. The interconduit pit membrane thickness (T_PM_) was suggested to be more important to embolism resistance than bordered pit area and pit aperture area (Lens et al. 2011; Li et al. 2016a). Based on a three dimensional view of pit membranes, it has been suggested that pit membrane thickness is related to the number of pore constrictions within a single pore pathway (Kaack et al. 2019; Zhang et al. 2020). Since the most narrow pore constriction within a pore pathway determines embolism resistance, the likelihood that the smallest pore constriction is very narrow will increase with the number of pore constrictions. Although this mechanistic link between pit membrane thickness and embolism resistance needs further research, higher embolism resistance in roots was found to correspond to thicker interconduit pit membranes in this study. Significant difference in pit membrane thickness between roots and stems, however, was only found for *C. avellana*. In fact, thicker pit membranes in roots of *C. avellana* may explain its more negative *P*_50_ and *P*_88_ values, which were 0.75 and 1.13 MPa more negative than stems of this species. Although interconduit pit membrane thickness is repeatedly shown to be related to embolism resistance (Jansen et al. 2009, 2018; Lens et al. 2011; Li et al. 2016a), this relationship cannot be treated as very tight, and might be blurred by potential artefacts in measuring embolism and pit membrane thickness (Kotowska et al. 2020).

We also do not fully understand the functional significance of the electron density of pit membranes between roots and stems. Since it is known that OsO_4_ treatment results in visualisation of unsaturated fatty acid chains of lipids (Riemersa 1968), differences in electron density may reflect different concentrations of lipids associated with pit membranes, which are likely to affect air-seeding and embolism resistance (Jansen et al. 2018; Schenk et al. 2017, 2018; Yang et al. 2020). Besides the hypothesis that xylem sap lipids represent remnants of vessel element cytoplasm (Esau 1965; Esau et al. 1966), it is possible that a high amount of vessel-associated parenchyma cells may contribute to a higher production of xylem sap lipids in roots than in stems (Morris et al. 2018; Schenk et al. 2018). Although the amount of parenchyma tissue in the species studied was not quantified, roots were found to show higher tissue fractions of ray and axial parenchyma than stems (Morris et al. 2016b; Plavcováet al. 2019), and it is also possible that the wide vessels in roots are more surrounded by parenchyma cells then narrow vessels in stem xylem (Morris et al. 2017).

Could the proximity of the stem and root samples, which was 15 to 20 cm in the saplings tested, explain the similarity in embolism resistance between both organs? Although this distance appears to be rather small, the samples selected were clearly root and stem samples from small saplings, with clear differences in conduit diameter and interconduit wall thickness between stem and root xylem. Since most vessels in the species studied are much shorter than 20 cm, with a mean vessel length below 6 cm (data not shown), it is unlikely that a few, long vessels could interconnect the root and stem samples that were used for our measurements. Moreover, the lack of any significant difference in embolism resistance between thick, mature root segments from forest saplings and the terminal root networks from potted saplings indicate that the proximity of the xylem tissue does not affect xylem embolism resistance in *C. avellana*.

Roots exhibited a wider conduit diameter and thinner interconduit wall thickness than stems in the three species studied, which is in line with previous studies (e.g. Aloni 1987; Anfodillo et al. 2012). Smaller conduits and thicker intervessel walls are suggested to increase hydraulic safety and mechanical support (Corcuera et al. 2004; Hacke et al. 2001; Mauseth and Stevenson 2004; Plavcová et al. 2019), although there is also evidence that drought-induced embolism is not directly related to conduit size (e.g. Choat et al. 2016; Klepsch et al. 2018; Skelton et al. 2018; Wason et al. 2018). Since roots were equally or even more resistant to embolism than stems in this study, this may indicate that conduit diameter and intervessel wall thickness do not play a major, direct role in embolism resistance. Although intervessel wall thickness was suggested to be linked with pit membrane thickness (Jansen et al. 2009; Li et al. 2016a), this relationship could not be supported for stems and roots of *A. campestre* and *A. pseudoplatanus* (Table 1). No clear relationship between these two features has also been reported in other studies (Klepsch et al. 2018; Scholz et al. 2013b).

In conclusion, this study shows that embolism resistance is higher for roots than for stems of *A. pseudoplatanus*, while no difference was found between roots and stems of *A. campestre* and *C. avellana.* This finding was supported by data on the interconduit pit membrane thickness, and suggests that the hydraulic vulnerability segmentation hypothesis does not apply to roots and stems of the three temperate tree species studied. Moreover, thick root segments of *C. avellana* show a similar embolism resistance to intact root networks, indicating that the pneumatic method can be applied to non-terminal plant material. We also demonstrate that the pneumatron approach shows promising results for establishing a high-throughput platform on a large number of samples, especially when this approach is combined with stem psychrometers (Pereira et al. 2020).

## Supporting information

Supplementary Table S1-S3, and will be used for the link to the file on the preprint site

## Acknowledgements

We thank the Botanical Garden and the Electron Microscopy Section of Ulm University for technical support. Daniel Glöckler and Peter Zindl are acknowledged for assistance with collecting plant material. M.W. acknowledges financial support from the Guangxi Education Department (GED). Y.Z. acknowledges financial support from the China Scholarship Council (CSC). This research was financially supported by the National Natural Science Foundation of China (No. 31800205, 31560124), and Guangxi Natural Science Foundation Program (No. 2015GXNSFBA139113).

