## Supplementary Table S1-S3, and will be used for the link to the file on the preprint site for "Root xylem in three woody angiosperm species is not more vulnerable to embolism than stem xylem"

**Supplementary Table S1:** Summary of the slope (S), $P_{50}$, and $P_{88}$ values obtained from xylem vulnerability curves based on the manual or automated pneumatic method for different organs from three species.

| Species | Organ | Pneumatic method | S | $P_{50}$ (MPa) | $P_{88}$ (MPa) |
|---|---|---|---|---|---|
| *A. campestre* | Stem | Manual | 10.37 ± 0.89 | -4.21 ± 0.38 | -9.15 ± 0.74 |
| | Root | Manual | 165.25 ± 39.26 | -4.86 ± 0.84 | -6.49 ± 0.97 |
| *A. pseudoplatanus* | Stem | Manual | 36.84 ± 13.27 | -2.63 ± 0.24 | -4.51 ± 0.49 |
| | Root | Manual | 27.89 ± 5.28 | -4.17 ± 0.31 | -6.26 ± 0.32 |
| *C. avellana* | Stem | Manual | 53.74 ± 24.29 | -2.77 ± 0.22 | -4.36 ± 0.34 |
| | Root | Manual | 31.74 ± 8.37 | -3.52 ± 0.33 | -5.49 ± 0.33 |
| | Root segment | Manual | 24.54 ± 4.45 | -3.41 ± 0.30 | -5.57 ± 0.61 |
| | Root | Automated | 106.35 ± 82.57 | -3.30 ± 0.38 | -4.91 ± 1.22 |

The fitting equation for vulnerability curves is given by $PAD = 100 / (1 + \exp((S/25)(\Psi - P_{50})))$, where PAD represents the percentage of air discharged, S is the slope of the curve, and $P_{50}$ is the xylem water potential at 50% of air discharged. $P_{88}$ (i.e. the xylem water potential at 88% of air discharged) was calculated following: $P_{88} = -2 / (S/25) + P_{50}$. Values are means ± SE, n = 3 or 5.

**Supplementary Table 2:** List of the t-statistics (*t*), the degrees of freedom (*df*), and the significance values (*P*) of *t*-tests for $P_{50}$ and $P_{88}$ values in stems and intact roots of the three species studied.

| Species | *t*-Test $P_{50}$ | | | *t*-Test $P_{88}$ | | |
|---|---|---|---|---|---|---|
| | *t* | *df* | *P* | *t* | *df* | *P* |
| *A. campestre* | 0.702 | 6 | 0.509 | -2.182 | 6 | 0.072 |
| *A. pseudoplatanus* | 3.924 | 8 | **0.004** | 2.982 | 8 | **0.018** |
| *C. avellana* | 1.882 | 8 | 0.097 | 2.381 | 8 | **0.045** |

*P* values in bold show significant differences.

**Supplementary Table 3:** List of the t-statistics (*t*), the degrees of freedom (*df*), and the significance values (*P*) of *t*-tests for $P_{50}$ and $P_{88}$ values in roots of *C. avellana*.

| Items | *t*-Test $P_{50}$ | | | *t*-Test $P_{88}$ | | |
|---|---|---|---|---|---|---|
| | *t* | *df* | *P* | *t* | *df* | *P* |
| Between I and II | -0.225 | 6 | 0.830 | 0.123 | 6 | 0.906 |
| Between I and III | -0.435 | 6 | 0.679 | 0.142 | 6 | 0.579 |

I = data of root networks based on the manual pneumatic method; II = data of thick root segments based on the manual pneumatic method; III = data of root networks based on the automated pneumatron method.
